# Defensive symbiont genotype distributions are linked to parasitoid attack networks

**DOI:** 10.1101/2024.07.24.604610

**Authors:** Taoping Wu, Anoushka A. Rodrigues, Tom Fayle, Lee M. Henry

## Abstract

Facultative symbionts are widespread in arthropods and can provide important services such as protection from natural enemies. Yet what shapes associations with defensive symbionts in nature remains unclear. Two hypotheses suggest that either interactions with antagonists, or host plants, may explain the prevalence of symbionts through shared selective pressures and routes of horizontal transmission. Here we investigate the factors driving similarities in the *Hamiltonella defensa* symbiosis shared among host species within field collected aphid communities. We show that, *Hamiltonella’s* genotype distribution strongly aligns with sharing the same parasitoids, rather than host plants, highlighting parasitoids as a key selective agent shaping the symbiosis across host species. Our data indicates parasitoid host-specificity drives the prevalence of specific aphid-*Hamiltonella* associations, suggesting defensive symbioses are maintained by the selective pressure imposed by dominant parasitoid species. These findings underscore the importance of interactions with natural enemies in explaining patterns of defensive symbiosis in nature.

## INTRODUCTION

Heritable bacterial symbionts are widespread in insects and often confer ecologically important traits to their hosts. Many insects harbour vertically transmitted obligate symbionts that have enabled expansions into novel feeding niches by provisioning their hosts with essential nutrients (Cornwallis *et al*. 2023; Jackson *et al*. 2022). Even more widespread are heritable facultative symbionts, which are not essential for hosts but can provide important context-specific benefits, such as protection from biotic or abiotic stresses (Heyworth *et al*. 2020; McLean & Godfray 2015; Tougeron & Iltis 2022; Wu *et al*. 2022). Facultative symbionts can also be horizontally transferred between hosts (Henry *et al*. 2015; Russell *et al*. 2003), and tend to be non-randomly distributed among plant-adapted insect populations and species (e.g. Henry *et al*. 2015; Henry *et al*. 2022; Jackson *et al*. 2023; Toju & Fukatsu 2011; Wu *et al*. 2022), although the reasons for this are currently unknown. This has led researchers to propose that facultative symbionts may provide benefits to populations in certain ecological niches, with high frequencies of symbiont carriage due to the acquisition and spread of symbionts that confer local adaptations (Jaenike 2012). However, the role of facultative symbionts in insect ecology has been hampered by our limited understanding of symbiont exchange networks and the agents of selection shaping their populations.

Insect facultative symbionts are often associated with providing protection against natural enemies (e.g. Oliver *et al*. 2003; Vorburger *et al*. 2010; Xie *et al*. 2014; Oliver & Perlman 2020; Zhao *et al*. 2023). While laboratory studies have provided valuable insight into defensive symbioses, our understanding of their network ecology in the wild is limited. Defensive symbionts are often maintained at intermediate frequencies in host populations (e.g. Ferrari *et al*. 2012; Osaka *et al*. 2010; Henry *et al*. 2013, 2015), suggesting a range of factors influence their spread and maintenance (Oliver *et al*. 2008; Smith *et al*. 2021). Selective forces such as the cost-benefit balance of maintaining symbionts in different ecological scenarios, as well as non-selective factors such as transmission rates, migration, and drift likely shape defensive symbiont distributions in host populations (reviewed in Vorburger 2022). Moreover, the horizontal transfer of facultative symbionts among host species complicates our understanding of how defensive symbioses function within insect communities, particularly those facing diverse natural enemies. The function of defensive symbioses is likely to have important implications for the dynamics of both natural ecosystems and agricultural landscapes, in which aphids are an important pest group (e.g. Leclair *et al*. 2021)

Some of the best-known examples of defensive symbioses have emerged from the bacterial symbionts of aphids. In addition to their obligate nutritional symbiont, *Buchnera aphidicola*, aphids harbour an array of facultative symbiont species, several of which can confer protection against natural enemies (Ferrari *et al*. 2004; Oliver *et al*. 2003). Among these, *Hamiltonella defensa*, hereafter *Hamiltonella*, stands out for its widespread presence in aphid populations and its ability to protect aphids from parasitoid wasp attack. In nature, aphid species and plant-adapted aphid ‘biotypes’, tend to be associated with only a few closely related *Hamiltonella* strains (Henry *et al*. 2013; Wu *et al*. 2022), however the reason for this is currently unclear. *Hamiltonella* is strongly associated with aphids that feed on certain host plants, suggesting interactions with plants may shape its distribution (Gimmi *et al*. 2024; Henry *et al*. 2013, 2015). However, different strains of *Hamiltonella* can also protect against different species of parasitoid wasps (McLean & Godfray 2015). In nature, some aphids have retained *Hamiltonella* strains that provide strong protection against their main parasitoid species (Gimmi & Vorburger 2024; Wu *et al*. 2022), suggesting antagonists may contribute to the symbiont’s distribution in nature. In addition, phylogenetic studies have shown that *Hamiltonella* can horizontally transfers between unrelated aphid species (Henry *et al*. 2013; Wu *et al*. 2022), although how transfer occurs in nature is unclear. Laboratory studies have shown that *Hamiltonella* can be transferred between hosts by parasitoids through a contaminated ovipositor (Gehrer & Vorburger 2012), but also via plant phloem (Li *et al*. 2018), suggesting that both mechanisms may contribute to the dynamics of symbiont acquisition and transmission in nature. However, the relative importance of plant and parasitoid interactions in shaping *Hamiltonella*’s presence in aphid communities remains unclear.

In this study, we analyse data from 1257 aphid samples to investigate ecological correlates linked to *Hamiltonella*’s distribution. Specifically, we ask whether *Hamiltonella* strains found in different aphid species are best explained by sharing parasitoids or host plants to shed light on the drivers of defensive symbiont dynamics in nature.

## MATERIALS AND METHODS

### Aphid-parasitoid sampling and identification

In 2021 and 2022, live and ‘mummified’ aphids were collected from 17 plant species in 26 locations across Greater London (Tables S1, S2), focusing on aphid species known to harbour *Hamiltonella* (from Henry *et al*. 2015; Wu *et al*. 2022). Aphids were collected by beating plants over a tray or manually from leaves and stems. To avoid resampling the same aphid clone or mummies from the same parasitoid, collections from the same plant species were spaced at least 10 meters apart. Additionally, 241 plant-aphid-*Hamiltonella* samples from a previous study (Wu *et al*. 2022) were included to control for potential sampling biases due to only sampling parasitised aphids. For robustness, we analyzed both the full data set of 1257 individuals and a conservative data set of 1016 samples collected in 2021 and 2022 from mummified aphids.

Aphid-parasitoid associations were identified using two methods: 1) Illumina amplicon sequencing of collected mummies, and 2) culturing live aphids until mummies formed and sequencing the emerged wasps. Mummified aphids were preserved in 70% ethanol for Illumina amplicon sequencing of the Cytochrome Oxidase subunit I (COI) gene to identify both parasitoid and aphid species. Live aphids, morphologically identified to species, were kept at 15°C with a 16h:8h light cycle for two weeks on host plant cuttings to harvest additional parasitoids. Emerging parasitoids were isolated, COI barcoded, and Sanger sequenced at Source Bioscience Ltd (London, UK) for identification. Morphologically identified aphids were also confirmed by Sanger sequencing.

DNA extractions were performed using DNeasy Blood and Tissue Kits (QIAGEN, Venlo, Netherlands). DNA from mummies was normalized to 5 ng/μL. A 407 bp COI gene region was amplified and sequenced using CS1/CS2 tagged primers and a modified PCR protocol (Table S5). Tagged PCR products were sequenced at the London Genome Centre. We sequenced 1108 mummies in two Illumina MiSeq runs using paired-end 300 bp reads: 370 samples in 2021 with 30,743 reads per sample, and 738 samples in 2022 with 26,281 reads per sample. Sequencing data was analyzed using the DADA2 pipeline (Callahan *et al*. 2016), with reads trimmed for quality, deduplicated, and chimeric sequences removed. Species were identified by comparing COI sequences to GenBank using BLASTn. Sequences were aligned using MUSCLE aligner in MEGA11 (Tamura et al. 2021).

Out of 1108 sequenced mummies, 833 yielded COI data for both parasitoid and aphid, with 129 detecting *Hamiltonella* for MLST. An additional 183 parasitoid samples identified by Sanger sequencing brought the total to 1016 parasitoid-aphid-*Hamiltonella* samples. Transient associations, where single aphids were collected from unrecognised host plants, were removed from the data set. Non-mummified samples, and those where *Hamiltonella* was not detected, were scored as having ‘No Association’ for parasitoid or symbiont and removed from corresponding analyses. Combined, these data sets resulted in 1257 samples for tripartite analyses.

### Hamiltonella strain identification

Samples from 2021/2022 were screened for *Hamiltonella* using diagnostic PCR targeting the 16S rRNA gene (Niepoth *et al*. 2018) (Table S5). Positive samples were genotyped using Multilocus Sequence-Typing (MLST) of four bacterial housekeeping genes: accD, hrpA, murE, recJ (Degnan & Moran 2008) (Table S5). PCR products were Sanger-sequenced in one direction and aligned using MUSCLE aligner in MEGA11 (Tamura *et al*. 2021). *Hamiltonella* sequences from aphids collected between 2011 and 2019 (Wu *et al*. 2022) were included in the alignment. The four genes were concatenated for analyses.

### Clustering of operational taxonomic units

COI gene sequences exhibiting over 99% similarity were clustered into a single Operational Taxonomic Unit (OTU). Species identifications for OTUs were based on the closest matching sequences in GenBank, with those >99% similarity assigned morphospecies name (sp1, sp2 etc). Multiple parasitoid OTUs matching the same species were designated as separate lineages (e.g., ‘clade 1’, ‘clade 2’) in analyses. For *Hamiltonella*, each MLST type was considered a separate OTU and referred to as a ‘strain’.

### Phylogeny reconstruction

Phylogenetic trees for aphids (COI), parasitoids (COI), and *Hamiltonella* (MLST) were constructed using the Maximum Likelihood method on the ATGC bioinformatics platform, visualized and rooted using iTOL software. *Adelges cooleyi*, *Formica fusca*, and *Hamiltonella* strain MEAM1 of *Bemisia tabaci* were used as roots for aphid, parasitoid, and symbiont phylogenies, respectively. Bayesian Information Criterion (BIC) was used for model selection through Smart Model Selection (SMS). Trees were generated in PhyML 3.0 (Guindon *et al*. 2010) using 100 bootstrap replicates.

### Network visualizations and specialization tests

Network visualizations were conducted using R v4.2.2 in RStudio. The aphid-*Hamiltonella*-plant and aphid-*Hamiltonella*-parasitoid interactions (Fig. 2) were mapped using the ‘igraph’ package (Csárdi *et al*. 2024) with the Fruchterman-Reingold layout. Networks showed similar patterns when reduced to data collected in 2021 and 2022. Bipartite plots (Fig. 4) were constructed using the ‘plotweb’ command in the ‘bipartite’ package (Dormann *et al*. 2008), assigning *Hamiltonella* strains to parasitoids proportionally. *Hamiltonella* aphid interactions matrix was plotted using the ‘ape’ package (Paradis & Schliep 2019) as in Wu *et al*. (2022). Network specialization was tested using the H2’ index (Blüthgen *et al*. 2007) by compare observed H2’ values with those from the null models using 1000 randomizations.

### Statistical analysis

Statistical analyses were conducted in R v4.2.2. Relationships between *Hamiltonella* strain diversity and parasitoid or plant diversity were tested using a linear model in the ‘lme4’ package (Bates *et al*. 2015), visualised in ggplot2 (Hadley Wickham 2016) and ‘igraph’ (Csárdi *et al*. 2024). Data were rarefied (n=1000 randomisations) to account for differences in sample sizes using the ‘rrarefy’ function in ‘vegan’. Diversity indexes were calculated using custom scripts. Mantel tests compared dissimilarity matrices of parasitoid, plant, and *Hamiltonella* communities using the Bray-Curtis dissimilarity matrix in the ‘vegan’ package with 9999 permutations. Aphid species lacking data for either parasitoids or *Hamiltonella* strains were excluded from analyses to prevent missing data from impacting results. This ensured accurate interpretation of relationships among aphids, parasitoids, and *Hamiltonella* strains.

## RESULTS

### Charactering the parasitoid and plant associations of aphids

Our analysis of the 1257 samples identified 45 parasitoid species and 36 *Hamiltonella* strains associated with 31 aphid species collected from 62 species of plants (Fig. 1, Table S1 and Table S2). A phylogeny based on the parasitoid COI gene revealed that the parasitoid species belong to 40 species (5 sub-species) from 12 genera and 2 Hymenopteran families, the Braconidae and Chalcididae. However, the vast majority (43/45) of aphid parasitoids were Braconids (Fig. S1). Most parasitoid species (65%) were recovered from more than one aphid species. However, most parasitoids were primarily associated with a single aphid species or genus (although the host species varied between parasitoids) with only intermittent use of other aphid species (Fig. 1). The majority (14/22 species, 64%) of aphid species were attacked by a single dominant parasitoid that accounted for >75% of parasitism for each aphid species, and most aphid species (18/31 species, 58%) fed on a single dominant species of plant (although again, the dominant parasitoid and plant species varied between aphid species). There were also numerous cases where multiple aphid species share the same host plant or parasitoid species (Fig. 1, Table S1). This demonstrates that this is an appropriate system to test whether the sharing of plants or parasitoids is linked to the sharing of *Hamiltonella* strains across aphid species.

**Figure 1:**
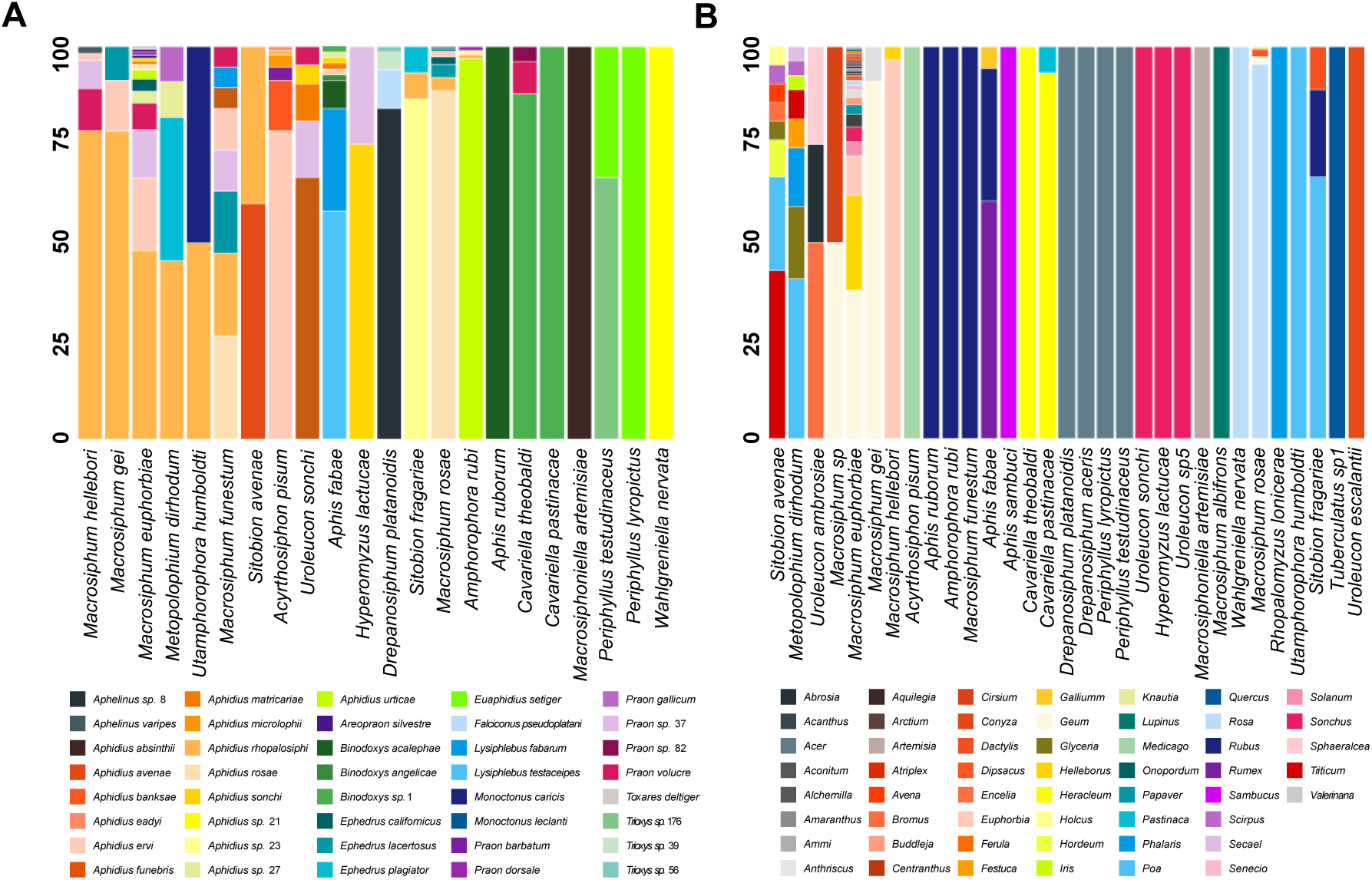
Relative frequency of parasitoid and host plant associations across aphids. Each bar represents an aphid species with different colours denoting the relative abundance of (A) different parasitoid species attacking the aphids and (B) different plant genera on which the aphids feed. Bars are organised according to the similarity of parasitoid/host plant communities as determined by Hierarchical Clustering. For visualisation purposes, different OTUs, or ‘clades’ belonging to a single parasitoid species were merged into a single colour.

### *Hamiltonella* diversity across aphid species

We were able to amplify *Hamiltonella* from 129 parasitoid mummies (Table S2). Genotyping the symbionts revealed 7 new *Hamiltonella* strains associated with aphids. By placing the new stains in a *Hamiltonella*-aphid phylogenetic matrix with previously published stains (Wu *et al*., 2022) we revealed that the newly identified strains tend to be associated with a single aphid species (Fig. S2). For example, *Hamiltonella* strain N_1_88 was only found in *Periphyllus lyropictus* and was found at relatively high frequencies (47%, 16 out of 34) in this aphid. However, we also found new cases of the same symbiont strains being shared by distantly related host species. For example, the aphids *Cavariella theobaldi* and *Macrosiphum funestrum* were both found with *Hamiltonella* strain N_2_65 and the aphids *Aphis fabae* and *M. funestrum* were both found harbouring strain N_4_75. In total, 15 out of 36 (42.6%) *Hamiltonella* strains were found across multiple aphid species. *Hamiltonella* strain 231 and strain 1485 are particularly noteworthy in their distributions; strain 231 was found in 14 aphid species across 8 genera and strain 1485 in six aphid species belonging to 3 genera. Our analysis also revealed several cases where a single aphid species was associated with two or more unrelated *Hamiltonella* symbiont strains. For example, *A. fabae*, *P. lyropictus*, *M. funestrum*, *Metopolophium dirhodum*, *Hyperomyzus lactucae*, and the *Medicago* biotype of *A. pisum* were all associated with multiple unrelated *Hamiltonella* strains that they carried at relatively high frequencies (Fig. S2).

We tested whether aphids associated with greater *Hamiltonella* diversity were associated with greater parasitoid or plant diversity using both rarefied and raw data sets (Table S3). Irrespective of the data set used, there was no correlation between *Hamiltonella* strain diversity, and the diversity of parasitoid or plant species associated with aphids (Fig. S3, Table 1).

**Table 1:** General linear model (GLM) of the association between *Hamiltonella* strain diversity and parasitoid/host plant diversity in aphids. Three data set were tested, including the full data set, rarefied (n = 7) data set and our conservative data set based on mummies only.

| Groups | Dataset | Richness |  |  | Shannon |  |  | Simpson |  |  |
| --- | --- | --- | --- | --- | --- | --- | --- | --- | --- | --- |
|  |  | R<sup>2</sup> | F-value | p-value | R<sup>2</sup> | F-value | p-value | R<sup>2</sup> | F-value | p-value |
| Parasitoid | Full dataset | 0.025 | 1.538 | 0.2293 | 0.0706 | 2.594 | 0.1229 | 0.041 | 1.899 | 0.1835 |
| Parasitoid | Rarefied n = 7 | -0.0369 | 0.5369 | 0.4778 | -0.0221 | 0.7194 | 0.4129 | -0.0258 | 0.6725 | 0.4282 |
| Parasitoid | Mummies only | -0.0683 | 0.0412 | 0.842 | 0.0001 | 1.002 | 0.3338 | 0.0052 | 1.078 | 0.3168 |
| Plant | Full dataset | 0.0121 | 1.368 | 0.2517 | 0.0249 | 1.767 | 0.1941 | 0.007 | 1.21 | 0.2803 |
| Plant | Rarefied n = 7 | -0.0563 | 0.0401 | 0.8437 | -0.0584 | 0.0074 | 0.9324 | -0.0584 | 0.0075 | 0.9322 |
| Plant | Mummies only | -0.043 | 0.3822 | 0.5463 | -0.0638 | 0.1001 | 0.7563 | -0.0645 | 0.0916 | 0.7666 |

### Aphid-*Hamiltonella* associations are correlated with parasitoid attack networks

Comparing Bray-Curtis indexes, we found that aphid species that shared the same parasitoid species tended to harbour the same *Hamiltonella* strains (Fig. 2A). However, aphid pairs that fed on the same species of plants had fewer incidences of sharing *Hamiltonella* strains (Fig. 2B). Using Mantel tests, we confirmed that there was a strong positive correlation between the similarity of *Hamiltonella* diversity shared by aphids and their parasitoid communities (Mantel test: r=0.294, p=0.0083**, Fig. 3A), but not their food plants (Mantel test: r=0.036, p=0.209, Fig. 3B). For robustness, we also analysed the data using only the samples collected in 2021 and 2022 and found the results were largely similar (Fig. S4 and S5).

**Figure 2:**
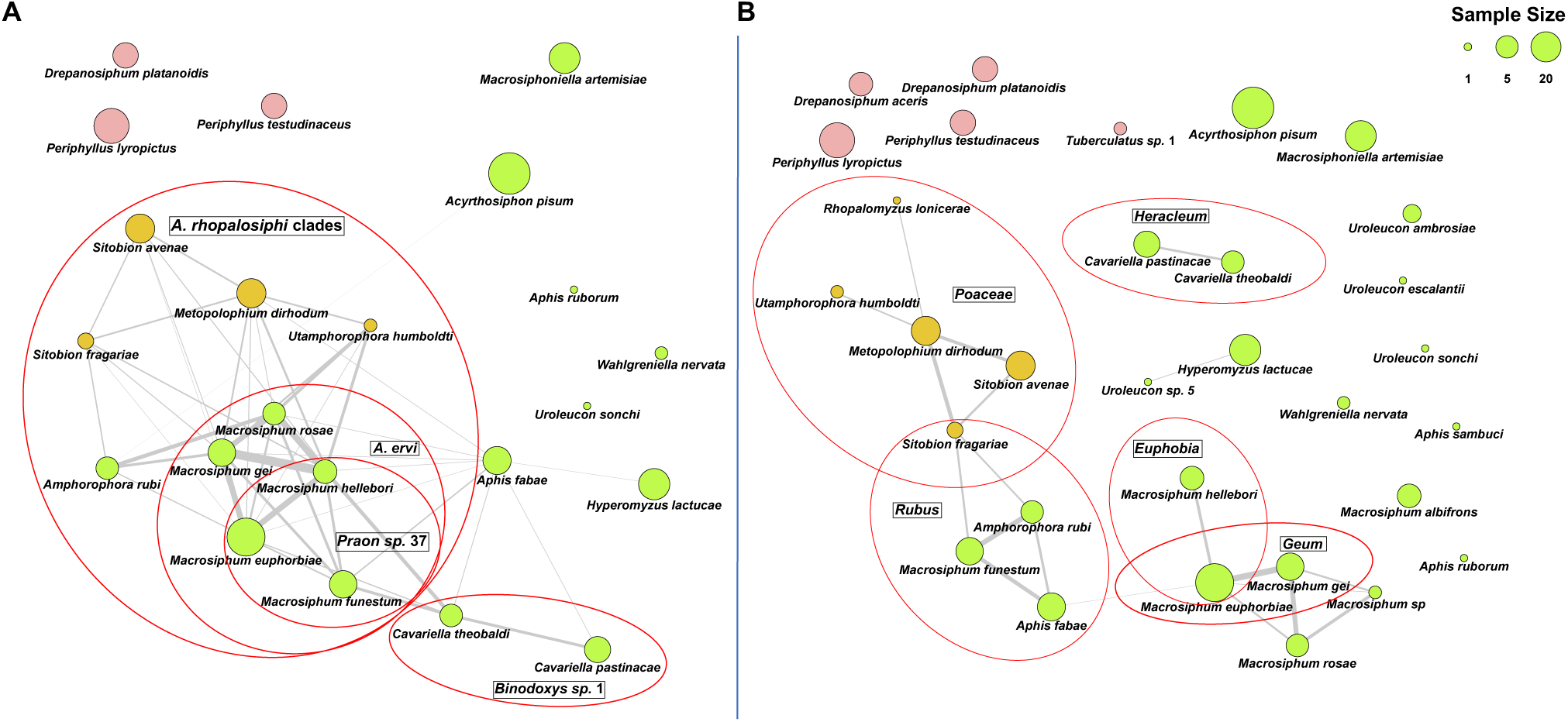
Parasitoid-*Hamiltonella* and plant-*Hamiltonella* network similarity of aphids. Grey lines connect aphid species that share similar *Hamiltonella* and (A) parasitoid species or (B) plant species. The thickness of each line corresponds to the sum of the Bray-Curtis similarity values of parasitoids/plants and *Hamiltonella* communities (Table S4). The size of each node reflects the number of *Hamiltonella* samples collected for each aphid species, while the colour denotes different aphid-plant classification: pink for tree-dwelling aphids, yellow for predominantly grass-dwelling aphids (although some host alternate), and green for herb-dwelling aphids. Red circles highlight predominant parasitoid and host plant associations (at > 0.1 Bray-Curtis similarity) that also share *Hamiltonella* strain(s). Parasitoid species and host plant genera (or families) are indicated in black squares.

**Figure 3:**
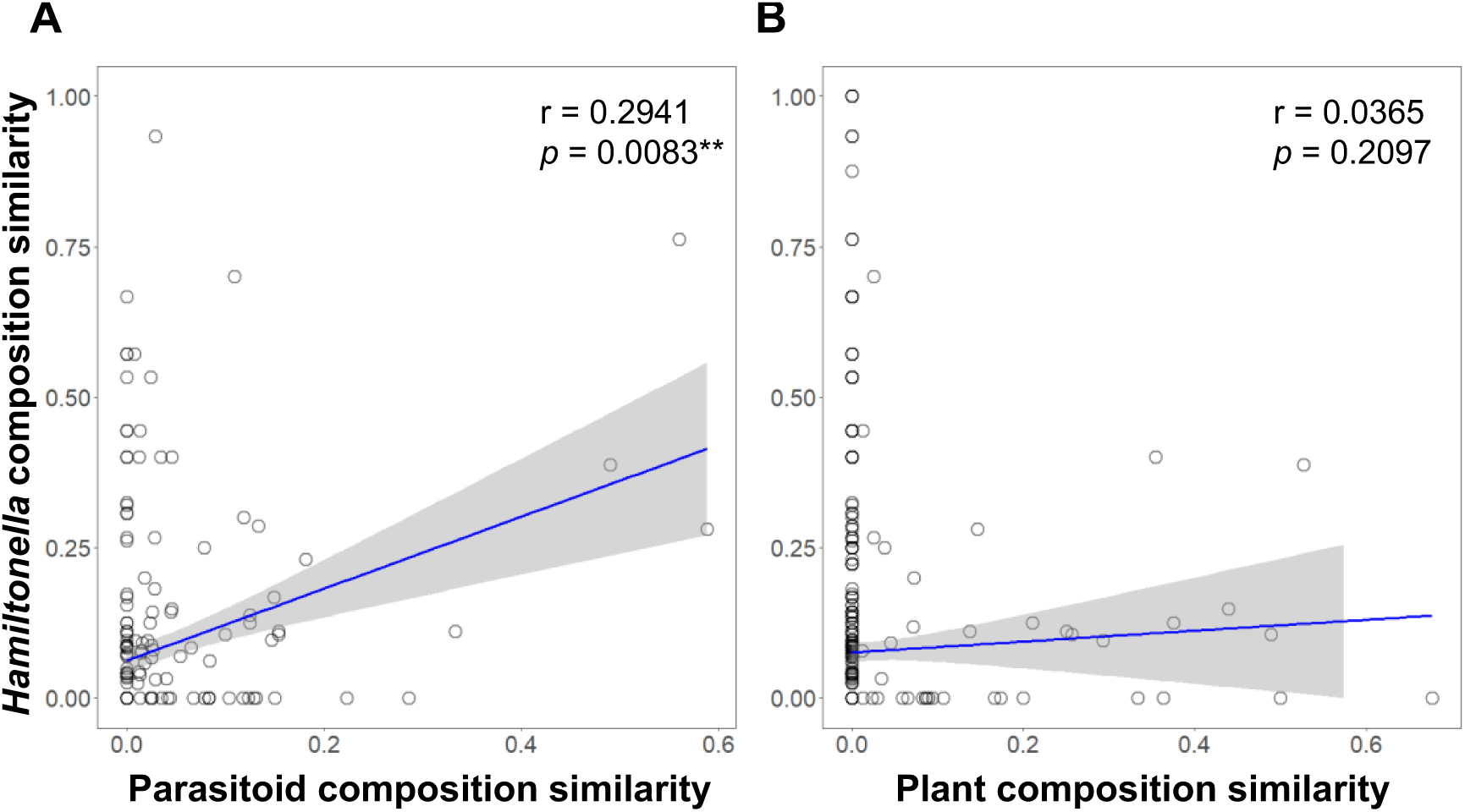
*Hamiltonella*-parasitoid and *Hamiltonella*-plant community similarity across pairs of aphid species. Each point represents the community similarities of a pair of aphid species. The x-axis represents the similarity in their (A) parasitoid, or (B) host plant community composition, while the y-axis represents the similarity in their *Hamiltonella* strain composition. Pearson correlation coefficients (r) and p-values (*p*) from Mantel tests are given in the upper right of each panel. Regression lines with 95% confidence intervals are plotted for visualisation purposed only, but not used in statistical analyses.

Within the parasitoid-aphid-*Hamiltonella* data, we identified 85 pairs of aphid species that shared *Hamiltonella* strains; among these, 47 pairs (55.3% of pair-wise comparisons) exhibited varying degrees of similarity in their parasitoid compositions (Table S4). 15 out of 19 (78.9%) aphid species sharing *Hamiltonella* strains could be linked by sharing at least one parasitoid species (Table S4). This is particularly evident in the *Macrosiphum* genus, which has the highest similarity in parasitoid species that attack them and tend to harbour the same *Hamiltonella* strains (average parasitoid Bray-Curtis similarity: 0.244, average *Hamiltonella* Bray-Curtis similarity: 0.423, Table S4). Several aphids that feed on grasses, including *Utamphorophora humboldti*, *M. dirhodum*, *Sitobion avenae* and *Sitobion fragariae* showed relatively high degree of parasitoids (Bray-Curtis = 0.156) and *Hamiltonella* similarity (Bray-Curtis = 0.074) with each other, as well as with *Macrosiphum* aphids (average parasitoid and *Hamiltonella* Bray-Curtis = 0.041 and 0.131, respectively, Table S4). The two *Caveriella* species, *C. theobaldi* and *C. pastinacae*, are also predominately attacked by the same parasitoid (Parasitoid Bray-Curtis = 0.33) and have evidence of sharing the same *Hamiltonella* strain 231 (Parasitoid Bray-Curtis = 0.11), but also feed on the same host plant, *Heracleum sphondylium*.

The plant-aphid-*Hamiltonella* data revealed 138 pairs of aphid species that shared symbiont strains; yet only 20 pairs (14.5% of pair-wise comparisons) had similarities in host plant species (Fig. 3B, Table S4). Furthermore, 8 aphid species, *A. pisum, Aphis ruborum, Aphis sambuci, Macrosiphum albifrons, P. lyropictus, Uroleucon escalantii, Uroleucon sonchi, Wahlgreniella nervata*, shared *Hamiltonella* strains with other aphid species but were not found sharing any host plants in our data set. Unlike the parasitoid-aphid-*Hamiltonella* interactions that linked all correlated aphid species together, the plant-aphid-*Hamiltonella* interaction formed three groups: i) 3 of 4 aphids feeding on *Rubus* are grouped with aphids that feed on grasses (Poaceae) through the host-plant alternating *Sitobion fragariae*, which are then weakly connected to 3 *Macrosiphum* species feeding on *Geum*; ii) 2 of 3 species on *Sonchus* share some similarity; and iii) the two *Cavariella* species that feed on *Heracleum* share plants, parasitoids, and *Hamiltonella* strains (Fig. 2).

Notably, there were 4 aphid species, *P. lyropictus*, *U. sonchi*, *W. nervata* and *A. ruborum*, that shared *Hamiltonella* strains but did not share the same parasitoids or host plants. This indicates that there were still connections in the symbionts’ distribution that cannot be explained by the plants or parasitoids identified in our data set.

### Specialised parasitoid-aphid-*Hamiltonella* associations

Our analyses revealed high degrees of specialisation in all interaction networks, indicating aphids, parasitoids, and symbiont strains tend to interact with a limited number of partners (Fig. 4). Specialization (as measured using H2’) was highest for parasitoid-aphid networks (Standardised Effect size (SES) = 153.3), but also high for both *Hamiltonella*-aphid (SES = 48.3) and the generated *Hamiltonella*-parasitoid (SES = 34.0) networks. Most aphid species harboured a single dominant *Hamiltonella* strain and were primarily targeted by a specific *parasitoid* species (Fig. 4A & 4B). For example, in the *Macrosiphum* genus, particularly *Macrosiphum gei*, *M. euphorbiae*, and *Macrosiphum hellebori*, all species predominantly harboured *Hamiltonella* strain 231 and were mainly attacked by a single parasitoid species, *Aphidius rhopalosiphi* clade 1. Notably, *A. rhopalosiphi* clade 1 was the most generalist parasitoid in our study, attacking 8 aphid species belonging to 4 genera; 6 of the species carried *Hamiltonella* strain 231. In addition to *A. rhopalosiphi* clade 1, *Macrosiphum* aphids were attacked by a similar group of parasitoids at lower incidences, including *A. rhopalosiphi* clades 2 and 3, and several species from the *Aphidius*, *Praon*, and *Ephedrus* genera (Fig. 4B). *Aphis fabae* also maintained diverse connections in that it shared *Hamiltonella* strains and parasitoids attacking it at relatively low frequencies with 10 other aphid species from 4 genera, including *Acyrthosiphon*, *Cavariella*, *Macrosiphum*, *Metopolophium*, and *Hyperomyzus* (Fig. 4, Table S1 & S5). Conversely, there were also aphid species, such as *Drepanosiphum platanoidis* and *Macrosiphoniella artemisiae*, that harboured a unique cluster of *Hamiltonella* strains and were each attacked either primarily or exclusively by a single parasitoid species. The parasitoid-*Hamiltonella* network also revealed several cases where parasitoids had particularly strong associations with certain symbiont strains, such as *A. rhopalosiphi* and *A. ervi*, which are linked to *Hamiltonella* strains 231 and 2578, respectively.

**Figure 4:**
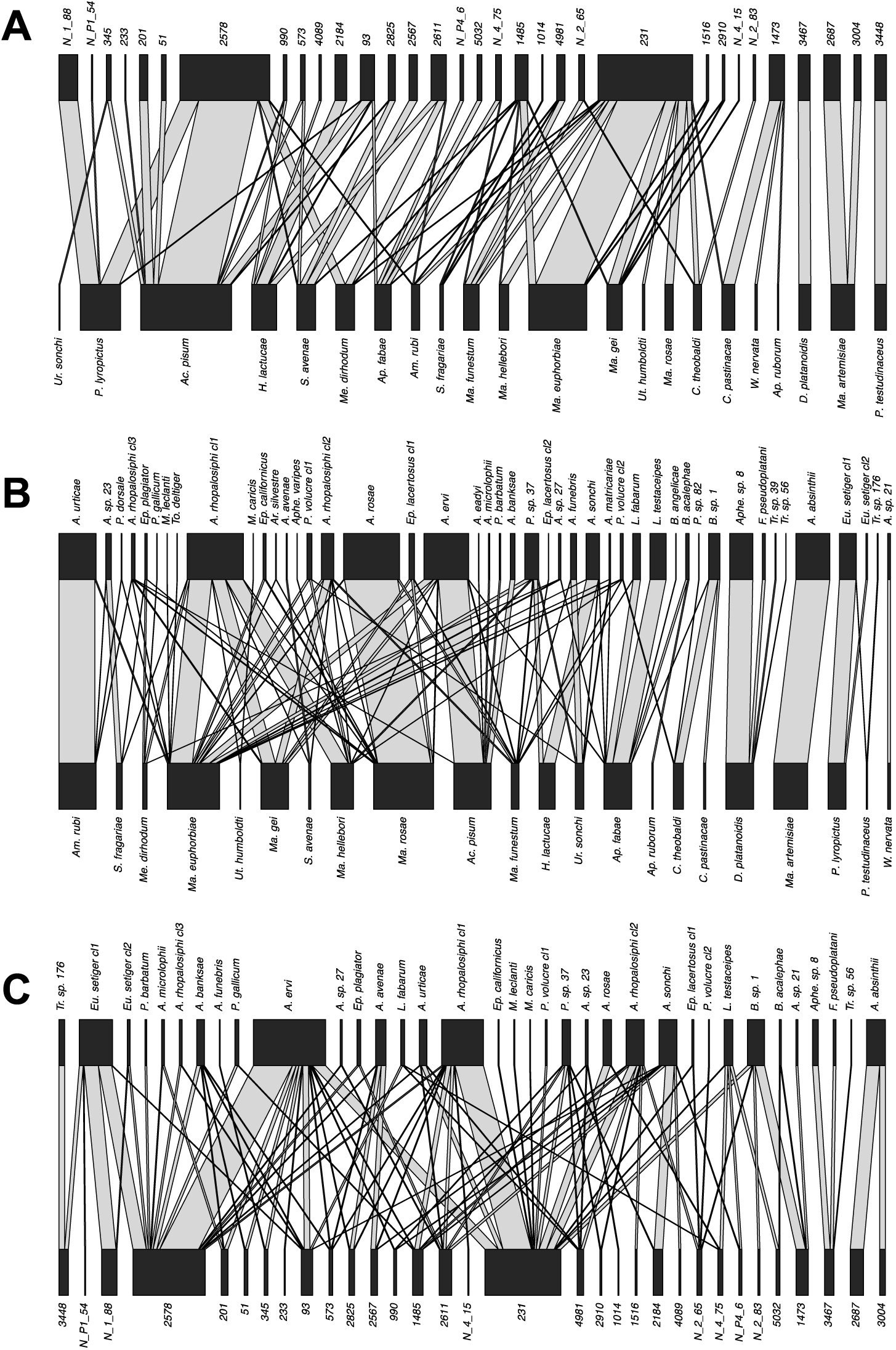
Bipartite parasitoid, aphid, and *Hamiltonella* interaction networks. Networks depict (A) *Hamiltonella* strain (top) – aphid species (bottom), (B) parasitoid species (top) – aphid species (bottom), and (C) parasitoid species (top) – *Hamiltonella* strain (bottom) interactions. The parasitoid-*Hamiltonella* network were generated by assigning *Hamiltonella* “individuals” in each aphid species to parasitoids attacking them, proportional to the frequency of attacks across different parasitoid species. Grey lines connect interacting species and strains, and the width of the line indicates the frequency of interaction between pairs. The width of the black bars denotes the relative abundance of individual aphids, parasitoids, and *Hamiltonella* stains used in each bipartite network, because these were assessed using different sets of samples.

## DISCUSSION

We demonstrate that natural enemy attack networks are linked to the distribution of defensive symbiont strains within an insect community. Population surveys, including ours, show that facultative symbionts providing protection tend to be non-randomly distributed across aphid species and plant-adapted populations; only certain species and populations harbour them, often with few symbiont strains at high frequency (Henry *et al*. 2013, 2015; Wu *et al*. 2022). We show aphids are typically attacked by a single dominant parasitoid species, and aphid species sharing the same parasitoids, rather than food plants, tend to carry the same strains of *Hamiltonella*. This suggests that interactions with parasitoids play a key role in the spread and maintenance of this defensive symbiosis within aphid communities.

### Parasitoids as selective agents shaping *Hamiltonella*’s distributions

Parasitoids impose strong selective pressure for the evolution of resistance in aphids. We show each aphid species is frequently attacked by a single dominant parasitoid, demonstrating a high degree of host specialization. We suggest parasitoid host-specificity may shape *Hamiltonella* distributions across aphid species. Laboratory studies have shown *Hamiltonella* strains vary in their degree of protection and specificity against different parasitoid species (Cayetano *et al*. 2015; Martinez *et al*. 2016). For example, McLean & Godfray (2015) found *Hamiltonella* strains associated with pea aphids on *Lotus* plants protect against *Aphelinus abdominalis*, but not *Aphidius ervi*, while strains on *Medicago* plants show the opposite trend. Frequent attack by a single parasitoid species likely explains why aphids carry a single or few closely related *Hamiltonella* strains, as these provide strong protection against their main natural enemy. Laboratory studies support this, showing attack by a single parasitoid species can lead to aphids carrying a single highly protective symbiont strain (Hafer-Hahmann & Vorburger 2020). Moreover, studies have found aphid species often carry *Hamiltonella* strains in nature that confer strong protection against their most common parasitoid (Gimmi & Vorburger 2021; Wu *et al*. 2022). Strikingly, we reveal that phylogenetically unrelated aphids attacked by the same parasitoids also carry the same *Hamiltonella* strains (based on 4 MLST genes). This suggests aphids retain similar symbiont strains to protect against a shared enemy. While the molecular mechanism of protection isn’t fully resolved, the genomes of most *Hamiltonella* contain a toxin-encoding bacteriophage known as APSE that is likely involved (Lynn-Bell *et al*. 2019). Variation in APSE’s toxin genes is thought to underlie the degree of protection against the parasitoid *Aphidius ervi* (Oliver & Higashi 2019). It would be in interesting to know whether APSE toxin variability also explains protection against different parasitoid species.

Although parasitoid and *Hamiltonella* community compositions were similar, there was no correlation between parasitoid species diversity and *Hamiltonella* strain diversity in aphids. Increased parasitoid diversity may not lead to increased symbiont diversity if there is a cost to resistance (Hafer-Hahmann & Vorburger 2024). Furthermore, factors such as transmission efficiency, host-symbiont compatibility, or interactions with other microbes, pathogens, or the environment, may also impact *Hamiltonella* diversity (Carpenter *et al*. 2021; Dykstra *et al*. 2014; Niepoth *et al*. 2018; Weldon *et al*. 2020; Goldstein *et al*. 2023). We also did not consider genotype diversity within parasitoid species, which may influence symbiont diversity associated with an aphid species (Hafer-Hahmann & Vorburger 2020).

### Influence of plants on *Hamiltonella*’s distribution

Studies show facultative symbionts occur more frequently in insect populations on certain plants, suggesting plant interactions may shape their distributions (Ferrari *et al*. 2004; Henry *et al*. 2013, 2015; Toju & Fukatsu 2011; Tsuchida *et al*. 2002). Examples include pea aphid biotypes on *Medicago sativa*, *Lotus pedunculatus*, and *Ononis* plants carrying biotype-specific *Hamiltonella* strains, and *A. fabae* populations feeding on different plants differing in *Hamiltonella* carriage (Gimmi *et al*. 2024; Henry *et al*. 2013, 2015). However, our results suggest there is no link between host plant sharing and the sharing of *Hamiltonella* strains in aphids. This indicates that plants have a limited role in shaping *Hamiltonella* distribution, at least among aphid species. Aphids on different plants might attract different parasitoid species due to changes in plant volatile profiles or *Hamiltonella* itself may modifying plant volatiles to attract different parasitoid species (Ahmed *et al*. 2022; Ali *et al*. 2022; Frago *et al*. 2017), leading to plant associated *Hamiltonella* strains. It would be interesting to determine if plant-adapted aphid biotypes are attacked by different parasitoid species, as this may explain their tendency to carry different *Hamiltonella* strains.

### Ecological vectors of *Hamiltonella* transmission

Hypotheses explaining horizontal transmission of facultative symbionts among insects include natural enemy and plant sap transmission. Natural enemies might pick up and transmit symbionts via contaminated mouthparts or ovipositors (Ahmed *et al*. 2015; Gehrer & Vorburger 2012; Kaech & Vorburger 2021; Soleimannejad *et al*. 2023; Tzuri et al. 2021), while plant sap transmission involves symbionts being released into plant sap and acquired by another insect ingesting it (Chrostek *et al*. 2017; Li *et al*. 2018; Pons *et al*. 2019). We show that the same *Hamiltonella* strains occur in distantly related aphids attacked by the same parasitoid species. Moreover, aphids carrying low frequencies of *Hamiltonella* strains typically found in other aphids, are often also attacked by their parasitoids at low incidences. This suggests parasitoids may be a vector of symbiont transmission in wild aphid populations. Laboratory studies have shown *Hamiltonella* can be transmitted by parasitoids in *A. fabae*, with transmission success depending on symbiont titre and haplotype (Kaech & Vorburger 2021). The likelihood of symbiont transfer can also be influenced by the relatedness of the hosts, and the symbionts they previously harboured (Łukasik *et al*. 2015; McLean *et al*. 2019). Parasitoids have also been shown to transfer facultative symbionts in other insects, such as *Myzus persicae* and *Bemisia tabaci*, and in house flies, *Musca domestica* (Ahmed *et al*. 2015; Soleimannejad *et al*. 2023; Tzuri *et al*. 2021). In contrast, we find little evidence that sharing the same host plants leads to sharing *Hamiltonella* strains. Plant-mediated horizontal transfer has been reported in several sap-feeding insects under laboratory conditions (Caspi-Fluger *et al*. 2012; Gonella *et al*. 2015; Li *et al*. 2017), including *Hamiltonella*, where plant transfer has been shown in *Sitobion miscanthi* feeding on wheat (Li *et al*. 2018). We find that when aphid species do share the same food plant and *Hamiltonella* strains, they also share the same parasitoids. Moreover, there are no cases where plant sharing alone explains *Hamiltonella*’s distribution, even in aphids that have a strong similarity in plant use, and do not share parasitoids. This suggests horizontal transfer by plants is either infrequent in aphids, or potentially only occurs in certain plants. It is also possible that selection from parasitoids may rapidly purge plant transferred symbiont strains from aphid populations, thereby contributing to the observed pattern. However, in most cases where *Hamiltonella* strains are shared between aphids it cannot be explained by plants, suggesting if it does occur it is of limited importance.

### Facultative symbionts as a reservoir of adaptations in insect defence

The non-random distribution of facultative symbionts in insects has puzzled scientists, especially in bacteria known to provide hosts with protection. Our results suggest that selection from natural enemies is key in shaping the distribution of defensive symbionts in aphids and potentially other insects. The link between *Hamiltonella* strains and parasitoid networks supports the idea that coevolutionary dynamics has led to host specialisation in natural enemies that has been largely mediated by the symbiont (Vorburger 2022). This may be due to symbiont-conferred defences being more specific than host-encoded defences, allowing hosts to tailor defences towards specific enemies through symbiont acquisitions. Specificity in symbiont-mediated protection has been reported in several defensive symbioses (Higashi *et al*. 2023; Łukasik *et al*. 2013; Mateos *et al*. 2016), and is particularly evident in *Hamiltonella* (McLean & Godfray 2015; Rouchet & Vorburger 2012; Wu *et al*. 2022). Bacteria have the potential to evolve more rapidly than the host’s genome, particularly when genes involved in protection are located on mobile genetic elements, such as the diverse toxin genes contained on *Hamiltonella*’s APSE phage. Parasitoids can become resistant to the presence of defensive symbionts (e.g. Dion *et al*. 2011; Oliver *et al*. 2008). The arsenal of defences carried by *Hamiltonella*’s APSE phages, in combination with their potential to mobilise, may be the crucial component in winning the evolutionary arms race against parasitoids, leading to the widespread distribution of the symbiont in aphids. However, the selection from *Hamiltonella* on specific natural enemies may also disrupt host specialization causing parasitoid to switch hosts to avoid highly protective symbiont strains, or provide an advantage to secondary parasitoids that attack a host at lower frequencies (e.g. McLean & Godfray 2017).

It has been suggested that horizontal transfer of defensive symbionts is rare enough that different host species carry distinct symbiont communities (Vorburger 2022). Our results suggest otherwise, as aphids attacked by the same parasitoid species tend to harbour related *Hamiltonella* strains. This suggests a more dynamic relationship where symbionts are maintained in host populations, at least across generations, at time scales that are relevant to selection from parasitoids. However, a finer-scale genetic analysis of shared symbionts and phages is needed to confirm this. Nonetheless, horizontal transfer of symbionts by parasitoids, even if rare, likely contribute to the genetic similarly of *Hamiltonella* strains occurring in aphids that share the same parasitoids and provides an important source of incoming symbiont variants for selection to act upon. Future studies are needed to assess whether symbiont transfer is rapid enough to counteract changes in parasitoid communities, or hamper parasitoid counteradaptations, to determine exactly how insects use symbionts in their defence against natural enemies.

## AUTHOR CONTRIBUTIONS

L.M.H. conceived the idea, acquired funding, led the investigation and supervision of the team. T. W. and A.A.R. conducted field collections, and formal analysis of the data. T.F. conducted additional analyses. All authors contributed to the drafting, reviewing, and editing of the manuscript.

All authors gave final approval for publication and agreed to be held accountable for the work performed therein.

## DATA ACCESSIBILITY

Data and scripts are available online: ########. The GenBank accession numbers for Sanger sequences determined in this study are ######## to ######## (*Hamiltonella* MLST). The GenBank accession numbers for Illumina sequences determined in this study are ######## to ######## (Aphid COI), and ######## – ######## (Parasitoid COI). The GenBank accession number for the raw data of Illumina sequencing used in this study is ##########.

## ACKNOLWEDGEMENTS

This work was funded by Leverhulme grant (RPG-2020-211) awarded to L.M.H.

